# TxNova: recovery of recurrent unannotated intergenic splice loci from existing bulk RNA-seq alignments

**DOI:** 10.64898/2026.08.24.746847

**Authors:** Zhao Li, Aaron W. James, Shengxuan Li

**Author notes:** Corresponding author: Zhao Li, M.D., Ph.D., 720 Rutland Avenue, Room 529, Baltimore, MD 21205,.

## Abstract

**Background:** Reference catalogs such as GENCODE capture most stably expressed mammalian genes but may not include condition-restricted or low-abundance transcripts. Reads supporting unannotated intergenic splice junctions are generally absent from annotation-restricted gene-count matrices; transcript assemblers may reconstruct a subset as novel models, but those models are typically handled separately from the annotated gene-count universe. TxNova is a lightweight command-line tool that directly indexes these unannotated splices from existing BAM files—without an external assembler, gffcompare, or workflow manager—yielding candidate leads for bench validation rather than standalone discovery claims.

**Methods:** TxNova retains CIGAR N junctions from STAR/HISAT2 BAM files that recur in ≥2 samples and are absent from a comprehensive annotation, clusters them into residual loci, and counts each locus alongside annotated genes in a unified matrix. Structure gates—canonical splice motif, same-strand distance, coverage valley, bridging-junction absence, minimum length—remove likely artifacts to yield structure-pass models. An optional contrast filter retains loci detected in treatment but nearly silent in control.

**Results:** A residual splice is a recurrent, unannotated CIGAR N junction that does not overlap any annotated gene body; intronic and antisense channels are out of scope. Across four published mouse treatment arms (GSE221720, GSE166522, GSE157460, GSE193335), harvest catalogs yielded 464, 657, 789, and 594 loci. On GSE221720, 45% of loci (209/464) shared an exact intron with another series, versus 0.074% for excluded junctions; a coordinate-placement control yielded 0/5,000 matches for length-matched intergenic decoys. Masked-gene recovery reached 87.6% (176/201) overall and 99.4% (176/177) among genes with a leak junction—sensitivity rather than precision estimates. An optional contrast provides a presence/absence screen on interval TPM. Residual models are partial reconstructions requiring cloning, RACE, or targeted proteomics for confirmation.

**Conclusions:** TxNova produces a reproducible intergenic residual-locus catalog from existing BAM files, with four downloadable mouse injury/infection catalogs. Cross-series recurrence and coordinate-based null controls support reproducibility above simple placement backgrounds but do not by themselves establish biological validity. The optional contrast step offers a practical screen for candidate leads warranting experimental validation.

## 1. Introduction

Reference annotations such as GENCODE serve as the canonical gene reference for most mouse and human RNA-seq studies ^1^. These annotations are large and carefully curated, yet may remain incomplete for condition-restricted or low-abundance transcription. Conditions such as bone fracture, implant infection, or inflammatory arthritis can produce spliced reads mapping to intervals that the annotation designates as intergenic. Such junction-supported signals are generally absent from annotation-restricted gene-count matrices; transcript assemblers may reconstruct some of them as novel models, but those models are handled separately from the annotated gene-count universe and can differ across samples. That transcription extends well beyond annotated gene boundaries is long established ^2^; what is notable here is that the sequencing data capturing these events have already been generated and aligned in many public studies.

Existing tools address related but distinct problems. Genome-guided assemblers reconstruct transcript models from alignments. Cufflinks ^3^, Scripture ^4^, StringTie ^5,6^, and Scallop ^7^ accept a BAM file and emit exon structures, aiming at a complete transcriptome: high recall of known isoforms at an acceptable novelty rate. Reference-guided StringTie (-G) uses the supplied annotation to guide reconstruction while still permitting novel transcript models ^5,6^. The resulting STRG/MSTRG models are therefore products of reference-guided transcript assembly rather than the output of a targeted intergenic splice census.

Meta-assemblers such as Cuffmerge ^3^ and TACO ^8^ merge per-sample GTF files. gffcompare ^9^ assigns class codes (u, x, i, j, …) against a reference. featureCounts ^10^ and stringtie -e -B ^5,6^ quantify a GTF. DESeq2 ^11^ and PyDESeq2 ^12^ test differential expression on a count matrix. CPAT ^13^, CPC2 ^14^, and Fickett’s TESTCODE ^15^ score coding potential. FEELnc ^16^ annotates long non-coding RNA. LeafCutter ^17^ clusters intron excisions to detect differential splicing, not novel intervals. Portcullis filters spurious splice junctions derived from alignments ^18^. PsiCLASS assembles transcripts from multiple samples simultaneously ^19^. ASJA extracts linear, back-splice, and fusion junctions ^20^. Large junction inventories such as intropolis document unannotated splices across public RNA-seq datasets ^21^. JunctionSeq tests differential usage of novel junctions at annotated genes ^22^. These junction-level resources catalog or test junctions without emitting locus models, whereas the assemblers emit unfiltered transcript models rather than an intergenic census (Section 3.2). Neither route yields intergenic locus models that are counted in the same matrix as GENCODE.

A comparable laboratory workflow can be assembled from StringTie, gffcompare, BEDTools, featureCounts, and CPAT joined by a workflow manager, but this composition introduces practical reconciliation steps. Strandedness conventions, contig naming, and GTF attribute versions must remain consistent across binaries; class-u transcript identifiers generated per sample must be reconciled across replicates; and short-read assembled isoforms are often partial reconstructions rather than full-length transcripts ^5,6^. TxNova packages the targeted residual-splice census, locus construction, counting, and filtering into one workflow while retaining the underlying BAM evidence.

TxNova indexes the splices that these pipelines leave behind. It accepts coordinate-sorted, indexed BAM files produced by STAR ^23^ or HISAT2 ^24^ and performs no alignment of its own. Gene, transcript, and exon records are imported from a comprehensive annotation (GENCODE comprehensive M39/GRCm39 for mouse, or a current GRCh38 release for human). If the species parameter is set to auto, the organism is inferred from the GTF; an explicit mouse or human value must agree with that inference. Only these two species are supported. The harvest step retains CIGAR N junctions that recur in at least two samples of the cohort yet are absent from the annotation, then clusters them into residual locus models that share a counting universe with annotated genes. Class assignment, fragment counts, coverage valleys, splice bridges, and coding scores are all computed on that universe directly from the BAM files.

A control-versus-treatment contrast is optional. When the sample sheet defines both groups, a secondary filter retains intervals detected in the treatment arm yet near-absent in the control arm, and PyDESeq2 may be run on the full locus matrix. When the sheet contains only a single group, or no group column at all, the pipeline still writes the residual catalog. The contrast filter is therefore a screen applied after discovery rather than the discovery step itself.

This paper makes three contributions. First, it formally defines a residual splice and implements TxNova as a unified command-line interface that recovers intergenic residual loci from existing STAR or HISAT2 BAM files. Second, it provides harvest catalogs for four published mouse treatment arms.

Third, it reports several validation checks on those catalogs, including masked-gene recall, cross-series intron recurrence with placement and failed-leak null distributions, and an optional contrast screen evaluated against a held-out label enumeration. The resulting models are partial reconstructions intended as leads for cloning, RACE, or targeted proteomics rather than as annotated genes.

## 2. Methods

### 2.1 Overview

TxNova is distributed as a single command-line entry point (txnova), driven by one YAML configuration file and one sample sheet. A Python orchestrator invokes an in-process Rust engine (txnova-core, via PyO3) for BAM, GTF, interval, and sequence operations. No workflow language is exposed to the user, and no external assembler, gffcompare, BEDTools, featureCounts, CPAT, or R/DESeq2 installation is required.

Each sample is supplied as a coordinate-sorted BAM file. The run annotation G is a GTF whose gene, transcript, and exon seqnames must form a subset of the BAM @SQ names, matched literally (chr1 versus 1 fails). The annotation must be comprehensive (GENCODE comprehensive, not basic), because a basic or cellranger-thin GTF causes annotated genes to appear intergenic. Preflight validation rejects mixed aligner families, mixed strandedness, truncated BGZF blocks, missing BAM indexes, and missing FASTA indexes. Only STAR and HISAT2 alignments are accepted; Bowtie2, minimap2, and GSNAP are rejected because their MAPQ scales are not interchangeable with those of STAR and HISAT2. At least two samples are required; the group column is optional.

The pipeline proceeds as follows: (1) write a slim copy of G (gene, transcript, and exon records only); (2) census CIGAR N junctions and retain cohort-recurrent junctions absent from G; (3) cluster those junctions into residual loci and write residual.gtf; (4) form the universe U = slim(G) ∪ residual models; (5) classify every transcript in U against G; (6) quantify every locus in U from every BAM; (7) scan structure features on intergenic representatives; (8) apply structure gates, and apply contrast abundance gates and optional differential expression only when both groups are present. Per-sample reconstructed GTF files are stubs; all novel models originate in the residual harvest. **Figure 1** summarizes this sequence.

**Figure 1.**
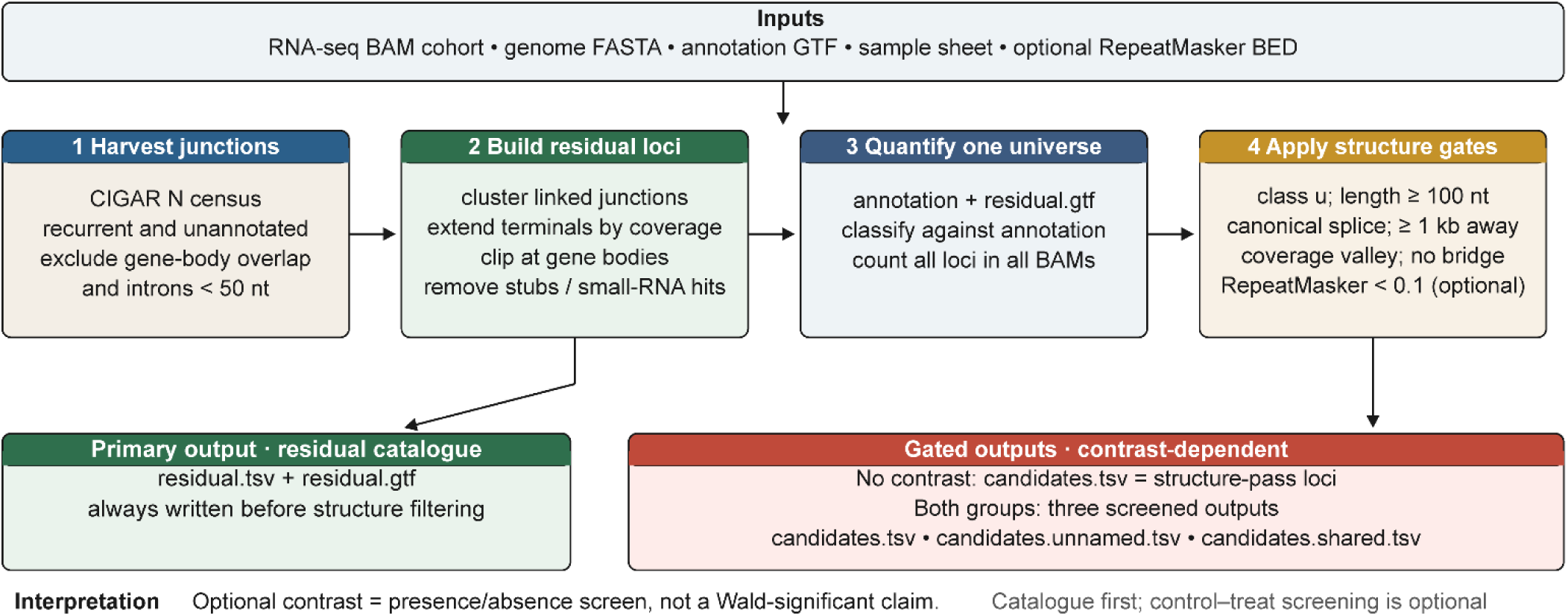
Residual-splice catalog and optional contrast screen. Junctions that are recurrent in the cohort and absent from the annotation are clustered into residual loci and counted alongside GENCODE in a single universe. Structure gates, including RepeatMasker filtering when a BED file is supplied, are applied when the gated candidate tables are written. A treatment-versus-control screen runs only when the sample sheet defines both groups. The four catalogs reported here are harvest residual tables; control BAM files were used only to label loci as silent or shared.

### 2.2 Residual splice harvest Junction census

For each BAM file, primary alignments that pass the MAPQ, duplicate, and (if requested) NH == 1 filters contribute CIGAR N operations. Deletions (D) are ignored, and paired-end mates are collapsed into a single fragment. The inferred transcript strand follows the library type (rf, fr, or unstranded).

A junction key is defined as (c, d, a, σ): contig, donor, acceptor, and strand. Let n_s,j be the fragment count of junction j in sample s. Cohort support comprises the sum of fragment counts and the number of samples with a nonzero count.

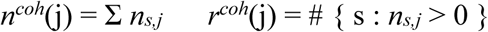

A junction is retained if it is cohort-recurrent (r_min = 2 by default, with at least two supporting fragments across the cohort) and is not an annotated intron of G. The harvest step itself ignores treatment and control labels.

When control samples are present, a junction is labeled silent if no control sample carries it, and shared otherwise; in the absence of control samples, the label is cohort. Shared junctions populate a separate structure-pass table when a contrast is defined and do not enter the treatment-detected final table.

A junction whose intron overlaps any gene body of G on either strand is discarded. Opposite-strand proximity without overlap is retained at this stage. Introns shorter than 50 nt are dropped ^25^.

#### Clustering into loci

Retained junctions on the same contig and strand are clustered using a union-find algorithm based on two rules, following the shared-site concept in LeafCutter ^17^ and the constitutive chaining approach used by Scripture-style assemblers ^4^: junctions share a donor or an acceptor, or their intron intervals overlap; alternatively, the open gap between adjacent introns is between 30 and 20,000 nt. This 30–20,000 nt range is an operational, configurable chaining window intended to accommodate plausible internal exons rather than a biological definition of exon size.

The representative intron chain is selected by cohort support and then by shorter introns. Terminal exons are extended by walking treatment coverage—or, where no treatment group is defined, coverage across all samples—outward from each splice site, up to 2 kb while tolerating gaps of ≤ 20 nt; an empty walk yields a 30 nt stub. Terminals that extend into a gene body are clipped, and loci retaining fewer than two exons after clipping are dropped.

Loci within 200 nt of a same-strand gene in G (or in an optional naming GTF) are removed during harvest as probable edge or read-through artifacts; this cutoff is evaluated on the intron span and is distinct from the 1 kb reporting gate applied later. Opposite-strand neighbors are not considered at this stage. Loci adjacent to small RNAs (snRNA, snoRNA, rRNA, tRNA, and related biotypes) on either strand are likewise removed; because small-RNA annotation is itself incomplete, this filter can both discard and retain true neighbors. The nearest_distance_bp column in residual.tsv reports the same-strand distance of the residual locus after terminal clipping (either-strand if the locus is unstranded), which represents the interval used by the 1 kb gate. Residual identifiers (RSDL.n) are assigned following a stable sort on junction count, cohort support, contig, and start position.

### 2.3 Universe, class, and quantification

The universe GTF comprises the slim annotation plus residual.gtf. Every transcript in U is classified against G using gffcompare-style codes ^9^. A locus is assigned class u only if all of its isoforms are class u, with no exon overlap and no gene-body overlap on either strand; the representative isoform is the longest class-u transcript. Counting is performed by fragment rather than by mate. Primary alignments with MAPQ ≥ 10 are used; NH uniqueness filtering is disabled by default; and duplicate skipping follows the BAM flag whenever the aligner has set it. A fragment is assigned to a locus only if every overlapping exon belongs to that single locus; fragments touching two loci are discarded (exclusive assignment at the locus level). Transcript counts within a locus are split equally, which has no effect on the locus-level values reported here. This scheme corresponds to neither featureCounts intersection nor union mode. Size factors for differential expression, when run, are computed from the full locus matrix.

### 2.4 Structure gates

These gates are applied when the gated candidate tables are written, regardless of whether a contrast is defined. The residual catalog (residual.tsv) reflects the harvest set and is written prior to gate application; the four treatment-arm catalogs reported in Results are harvest tables.

Canonical splice. Introns must exhibit GT-AG or GC-AG dinucleotides on the transcript strand (U2-type pairs ^26^). AT-AC (U12) junctions are not accepted. Default: zero non-canonical junctions. This is a structural-plausibility gate; a canonical motif alone does not establish biological validity.

Same-strand distance. A minimum distance of 1,000 bp to the nearest same-strand gene in the run annotation is required. The 1 kb value is a configurable operational intergenic buffer, consistent with conservative separation conventions used in lincRNA cataloging ^27,28^, rather than a biological boundary or an estimate of median gene-to-gene spacing in GENCODE M39. It does not imply that intergenic sequence is free of selective constraint at this scale ^29^.

Coverage valley. If a same-strand neighbor is present, at least one detecting sample must exhibit a local coverage dip together with a low intervening mean (default window: 50 bp; valley: 200 bp; mean depth ≤ 1 or ≤ 0.1 of locus depth). This criterion rejects transcribed 3′ tails that contain only a single empty window.

Bridging junction. The presence of two or more spliced fragments joining the locus to the nearest same-strand gene results in locus rejection.

Length. The minimum spliced length is 100 nt. GENCODE and lincRNA catalogs apply a 200 nt biotype floor ^27,30^; the lower default here retains two-exon residual models of 100–199 nt.

RepeatMasker. When genome.rmsk_bed is specified, RepeatMasker coverage of the spliced model is computed. The ≥ 0.1 discard threshold is a configurable heuristic used only when the gated candidate and contrast tables are written (Section 2.5), never for the harvest catalog. All four runs were configured with the mm39 RepeatMasker BED file. Section 3.4 reports this fraction on the unfiltered GSE221720 harvest set as a post hoc annotation (high, ≥ 0.1; low, < 0.1). Harvest deliberately retains repeat-rich models so that repeat-associated signal is not erased from the census; however, four-series shared hits are nonetheless dominated by that class. The discard threshold was applied only in the periodontitis contrast (Section 3.5).

### 2.5 Optional contrast filter

When the sample sheet contains at least one control and one treatment sample, structure-pass class-u loci are partitioned by heuristic abundance gates. These values are configurable screening parameters rather than universal expression cutoffs; their scale is informed by prior work on distinguishing low-level RNA-seq expression ^31^. candidates.tsv retains intervals with a control maximum TPM < 0.5, at least three treatment samples at TPM ≥ 0.1, a treatment median TPM ≥ 0.5, and passage of the RepeatMasker and structure gates. candidates.unnamed.tsv contains structure-pass loci with a control maximum TPM ≥ 0.5, and candidates.shared.tsv contains structure-pass loci carrying at least one harvest junction labeled as shared.

In the absence of both groups, candidates.tsv simply comprises the structure-pass residual set. The contrast filter operates on the presence and absence of interval TPM; with four treatment and four control samples, a three-of-four versus zero-of-four pattern corresponds to a one-sided Fisher exact P ≈ 0.07, and DESeq2 assigns such rows the low_count label in nearly all cases (Section 2.6). Loci with shared harvest junctions are excluded from the final contrast table.

### 2.6 Differential expression

PyDESeq2 ^12^ is optional. For residual rows in this paper, it serves as a size-factor annotation: independent filtering leaves padj undefined (low_count). It is not the mechanism by which residual loci are identified.

Default labels for a contrast row that has already passed the hard gates are wald (padj defined using Benjamini–Hochberg adjustment ^32^, padj < 0.05, log2 fold change ≥ 0.5) or low_count (padj NA, Wald P exists, same fold-change cutoff). The log2 fold-change threshold is a workflow setting rather than a criterion defined by the Benjamini–Hochberg procedure. All residual rows reported below are labeled low_count.

### 2.7 Coding scores and rank

To assist users in judging whether a recovered interval appears non-coding (lncRNA-like) or carries a candidate short open reading frame (an sORF/micropeptide lead), TxNova scores the best ORF of each locus using a packaged hexamer-based coding score, defined below. The representative sequence is spliced (with negative-strand models reverse-complemented) and scanned in three frames for ATG…TAG/TAA/TGA; the longest ORF is retained, with a configurable default minimum reported length of 50 aa and require_orf disabled. Short ORFs can encode functional peptides ^33^, but 50 aa is an operational reporting threshold rather than a biological definition of an sORF. In-frame hexamers of that ORF are scored against the packaged CPAT hexamer tables for the resolved species ^13^ using a Laplace floor of 10⁻⁸. The statistic is a hexamer log-likelihood ratio, not CPAT’s logistic model. It was evaluated on a reservoir sample of GENCODE M39 transcripts (2,000 protein-coding and 2,000 lncRNA; seed 20260818; 1,999 and 1,997 scored), yielding an area under the receiver operating characteristic curve (AUC) of 0.90. At the sign threshold of 0, sensitivity is 95.4% (1,908/1,999) and specificity is 47.6% (951/1,997); Youden’s J is maximized at 0.21 (sensitivity 80.3%, specificity 85.9%). Threshold 0 is therefore a high-recall inspection flag rather than a calibrated coding-versus-lncRNA cutoff. The product default remains 0; the column is not used as a discovery gate; and Fickett TESTCODE ^15^ is reported separately.

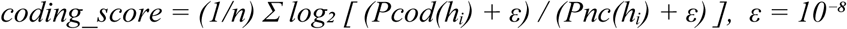

Software inspection labels are: no ORF ⇒ noncoding; score > 0 ⇒ coding; score < 0 ⇒ noncoding; score = 0 ⇒ ambiguous. These labels are heuristic inspection categories tied to the product threshold and do not constitute biological coding/noncoding calls. Structure-pass loci are ranked for inspection. The rank represents an inspection order, not a gene call.

### 2.8 Implementation

BAM I/O uses rust-htslib, a Rust binding to htslib ^34^. Concurrent sample workers scale according to available RAM (approximately 2 GiB per worker plus a 4 GiB reserve). Configuration is handled by Pydantic v2 with extra=“forbid”. Every gate threshold is exposed as a YAML key. Stamps skip unchanged stages. TxNova is installed via pip install txnova (Linux and macOS wheels; Windows via WSL2). The version described and used herein is 0.1.8 (https://doi.org/10.5281/zenodo.21970482).

Source and documentation: https://github.com/leelieber2025/TxNova, https://txnova.readthedocs.io/

### 2.9 Held-out gene recall

From genes with a mean count ≥ 1 and at least two exons in the GSE221720 4v4 design (n = 22,604), 1,000 gene_ids were drawn with seed 20260818 and removed from both the harvest GTF and the naming GTF. Isolated genes were defined as those whose body lies ≥ 1 kb from any remaining same-strand gene, with no overlap on either strand (n = 201). A leak hit is a recurrent unassembled CIGAR N junction whose intron overlaps a hidden gene, and residual recall is defined as a residual locus overlapping that gene. Conditional recall, computed among isolated genes that still produced a leak junction, is 176/177 (99.4%); unconditional recall of the full isolated set is 176/201 (87.6%). That run produced 1,741 residual loci, because hiding 1,000 annotated genes causes those gene bodies to appear intergenic; 1,741 is therefore not comparable to the 464 loci of the unmasked run. Separately, 5,000 annotated protein-coding introns were relocated into intergenic sequence, preserving intron length and a GT-AG or GC-AG pair while remaining ≥ 200 nt from any gene body (seed 20260818). A decoy was counted as a hit only if its coordinates coincided with a treatment-recurrent unassembled junction in the unmasked GSE221720 leak table. This constitutes a placement control rather than a BAM-injection test.

## 3. Results

### 3.1 Four treatment-arm series

Residual catalogs were generated from the treatment arm of four published mouse bulk RNA-seq series, realigned with STAR to GRCm39 and counted against GENCODE M39 comprehensive rather than the basic GTF (**Table 1**). Junction inclusion required recurrence in at least two treatment samples. Control alignments, where present in the sample sheet, were used only to label each locus as silent or shared. GSE221720 was originally aligned with HISAT2 ^35^; all catalogs and the StringTie comparison reported below utilized the same STAR BAM files.

**Table 1.** Treatment-arm series used for residual catalog generation.

| Accession | Treat arm | n | Library |
| --- | --- | --- | --- |
| GSE221720 | ligature periodontitis bone <sup>35</sup> | 4 | polyA, rf |
| GSE166522 | implant osteomyelitis, day 3 <sup>36</sup> | 3 | polyA, unstranded |
| GSE157460 | ulnar stress fracture, day 1 <sup>37</sup> | 6 | Ribo-Zero total RNA, rf |
| GSE193335 | collagen-induced arthritis, week 6 <sup>38</sup> | 5 | polyA, rf |

Each original study reported differential expression for annotated genes. The same BAM files also contain unannotated splices. Contralateral controls are included for GSE221720 alone in Section 3.5.

### 3.2 StringTie class u versus the residual census

Reference-guided StringTie uses the supplied annotation to guide transcript reconstruction while still permitting novel models ^5,6^. In these datasets, the resulting class-u/MSTRG models provide an assembly-based comparator rather than a targeted intergenic splice census. TxNova instead records recurrent extra-genic junctions directly before locus construction. The comparison employed StringTie 3.0.3 on the same STAR BAM files and GENCODE M39 GTF: stringtie BAM -G annotation.gtf -o sample.gtf -p N per sample, with --rf for stranded libraries, followed by stringtie --merge -G annotation.gtf. The GSE221720 merged GTF contains 491,372 transcripts because merge -G writes the entire GENCODE M39 comprehensive set (481,956 transcripts) alongside leftover isoforms; 454 loci are all-u, of which 395 are MSTRG. After the same structure and detection gates, 7 MSTRG loci remained on GSE221720 and none entered the final contrast table. All three comparisons had both groups defined in the sample sheet. On a single treatment BAM (bl_2), StringTie with -G wrote 12,087 per-sample transcripts lacking a reference_id, and 35,469 STRG transcripts without -G. The corresponding splice census was not empty: 13,088, 19,198, and 22,877 treatment-recurrent CIGAR N junctions had no counterpart intron in the StringTie merge (**Table 2**). That set is distinct from the TxNova leak table (94,348 treatment-recurrent unassembled junction records on GSE221720 against GENCODE; Section 3.3). Most of those 94,348 records overlap a gene body and lie outside the intergenic residual channel by design.

**Table 2.** StringTie 3.0.3 reference-guided merge versus the residual census on three treatment arms. All-u loci, MSTRG u, MSTRG after gates, and MSTRG final are class assignments against GENCODE derived from the StringTie merge, independent of the TxNova harvest engine. Unassembled N is counted directly from BAM CIGAR N operations against the same merge and is likewise decoupled from the harvest pipeline (script archived with the release).

| Series | MSTRG u | After gates | Final | Unassembled N |
| --- | --- | --- | --- | --- |
| GSE221720 | 395 | 7 | 0 | 13,088 |
| GSE166522 | 147 | 4 | 0 | 19,198 |
| GSE157460 | 1,007 | 9 | 0 | 22,877 |
| GSE193335 | 1,424 | 0 | 0 | 3,955 |

### 3.3 Residual catalogs (no contrast)

Junctions present in at least two treatment samples and absent from GENCODE M39 were clustered. Harvest excludes gene-body overlap, same-strand neighbors within 200 nt, and small-RNA neighbors. Structure gates (canonical splice, 1 kb distance, valley, bridge, and 100 nt length) together with the RepeatMasker 0.1 discard are applied when the gated candidate tables are written (Sections 2.4–2.5). Structure-pass counts are 247, 204, 186, and 255. Control alignments labeled each locus as silent or shared without affecting catalog membership.

On GSE221720, 94,348 treatment-recurrent unassembled CIGAR N junction records were observed, corresponding to 94,083 unique coordinates, because some junctions recur at the same coordinate across the sample set. By record count, 93,451 overlap a gene body on either strand and 897 are intergenic. Of those 897 intergenic records, 626 became residual-locus introns (464 loci) and 271 failed the 200 nt cutoff, the small-RNA filter, clustering, or the two-exon clip. The failed-leak set used in Section 3.4 is coordinate-based throughout: the 94,083 unique unassembled coordinates, minus the 626 that are residual-locus introns, yield 93,457 failed-leak coordinates. The 13,088 junctions in Table 2 are treatment-recurrent N operations missing from the StringTie merge, a different reference computed independently of this harvest table. Sequential structure-gate losses, applied in the order given and without RepeatMasker, are as follows: GSE221720 length 0, canonical 20, 1 kb 97, valley 99, bridge 1 (247 remain); GSE166522 6, 19, 305, 119, 4 (204); GSE157460 4, 441, 66, 92, 0 (186); and GSE193335 30, 205, 40, 63, 1 (255). The canonical-splice loss on GSE166522 (19/657, 2.9%) is consistent with the other polyA series because the unstranded intron scan tests both genomic readings rather than defaulting to the plus strand (Section 2.2); defaulting to a single reading inflates the apparent canonical-splice failure rate of an unstranded library by approximately eightfold. The Ribo-Zero series GSE157460 loses the most loci at that gate (441/789, 55.9%). Differences in library composition, including a greater contribution from precursor- or repeat-derived RNA in total-RNA data, are one possible explanation, but this was not tested directly here.

Shared harvest labels predominate (401/464, 591/657, 688/789, and 528/594); thus, most residual splices are not treatment-restricted. Median spliced lengths are 716, 825, 467, and 396 nt (**Figure 2**). On GSE221720, a 30 nt terminal stub occurs at 2/464 5′ ends and 1/464 3′ ends, and 317/464 3′ exons exceed 200 nt; consequently, the random-like polyadenylation signal (PAS) rate is not explained by empty walks. Because most loci are shared, the harvest catalog in Table 3 serves as a discovery-stage index rather than a candidate list: given matched control samples, the presence/absence contrast filter of Section 2.5 (demonstrated in Section 3.5) represents the intended route from that shared background to a small treatment-detected, control-silent screen.

**Figure 2.**
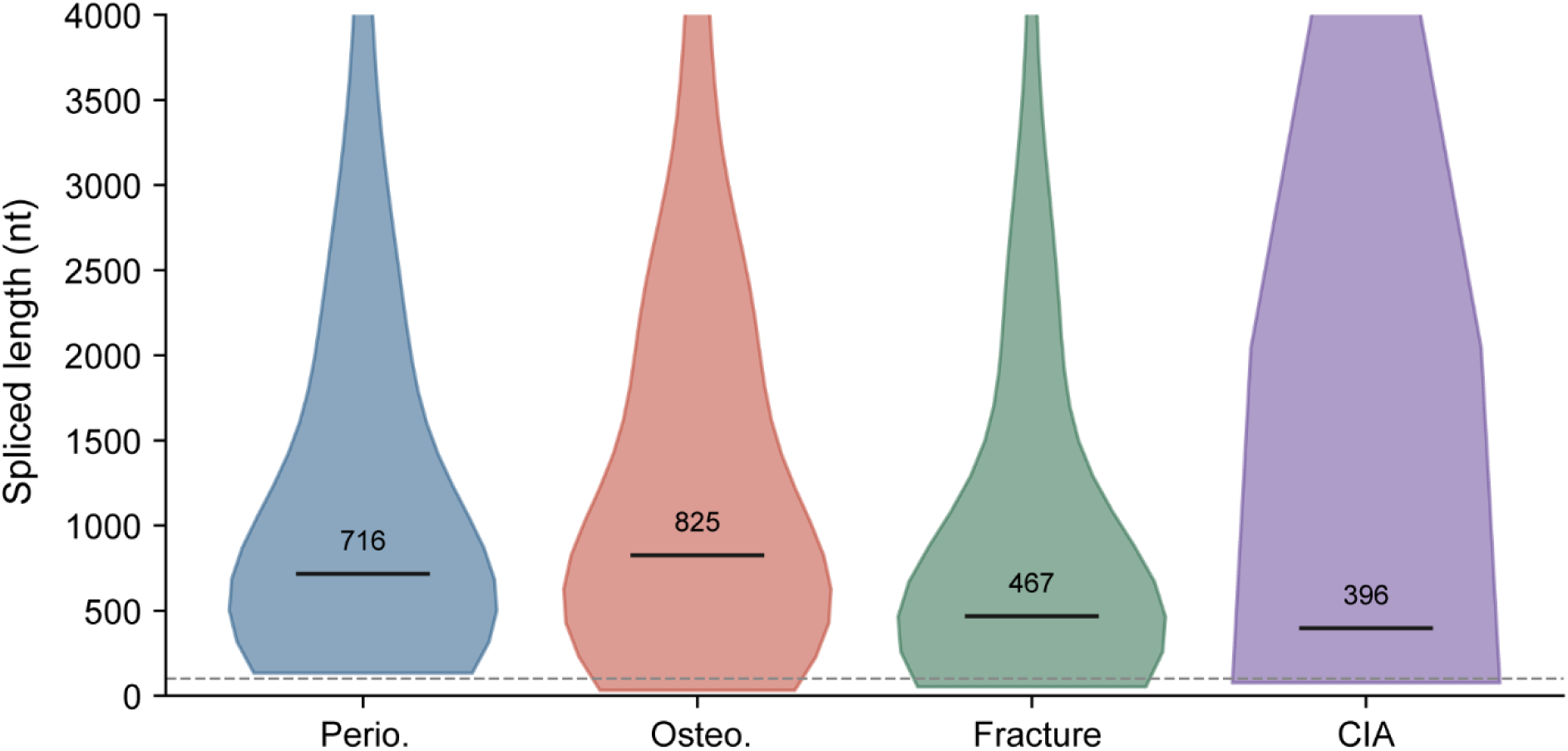
Spliced-length distribution of harvest residual loci across the four treatment arms. Vertical numbers indicate series medians (716, 825, 467, 396 nt). The dashed line denotes the 100 nt reporting floor. The vertical axis starts at 0 and is truncated at 4,000 nt.

**Table 3.** Harvest residual catalogs for four treatment arms. This represents the harvest set rather than a candidate-gene table. Silent and shared are harvest-junction labels and do not denote biological absence in the control arm. Structure-pass counts apply the canonical-splice, 1 kb distance, coverage-valley, no-bridge, and 100 nt criteria, without the RepeatMasker discard.

| Series | Residual loci | Silent | Shared | $\geq 2$ junctions | $\geq 3$ exons | structure-pass |
| --- | --- | --- | --- | --- | --- | --- |
| GSE221720 | 464 | 63 | 401 | 136 | 109 | 247 |
| GSE166522 | 657 | 66 | 591 | 204 | 154 | 204 |
| GSE157460 | 789 | 101 | 688 | 155 | 117 | 186 |
| GSE193335 | 594 | 66 | 528 | 109 | 88 | 255 |

Reporting thresholds alter catalog size (Figure 3). Increasing the same-strand distance from 200 bp to 5 kb, or the spliced-length floor from 50 nt to 500 nt, shrinks every series monotonically; requiring a harvest junction in at least three treatment samples (r_min = 3) leaves 80, 107, 168, and 91 loci. Harvest employed r_min = 2 and the 200 nt cutoff; the 1 kb and 100 nt values scanned in Figure 3 are reporting thresholds rather than filters already applied to Table 3.

**Figure 3.**
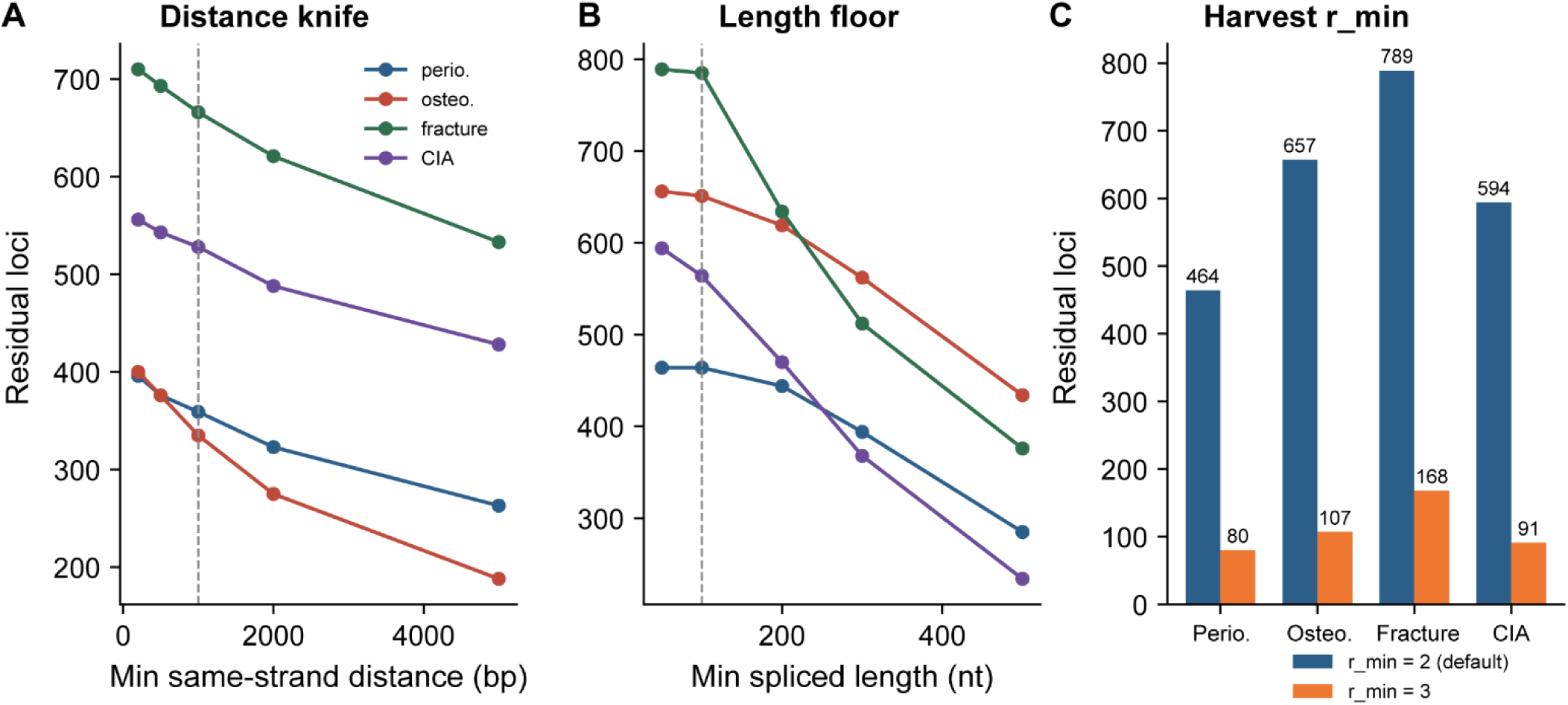
Catalog size as a function of reporting thresholds. (**A**) Minimum same-strand distance. (**B**) Minimum spliced length. (**C**) Harvest r_min = 2 versus r_min = 3. Dashed lines indicate the defaults.

After masking 1,000 randomly chosen expressed multi-exon genes (seed 20260818), two recoveries must be distinguished. Conditional sensitivity, among isolated genes that still produced a leak junction, is 176/177 (99.4%). Unconditional recovery of the isolated set, including genes the census never observed, is 176/201 (87.6%). Neither quantity estimates precision for residual intervals in the unmasked run. None of the 5,000 length-matched intergenic decoy coordinates coincided with a treatment-recurrent unassembled junction in the GSE221720 leak table (0/5,000), against 897 observed intergenic unassembled junctions, 626 of which entered residual loci. Under this exact-coordinate placement control, random intergenic GT-AG pairs of annotated intron length rarely coincide with the recurrent unassembled junctions being cataloged; because the decoys carry no simulated read support, this result does not estimate the false-positive rate of individual residual loci in Table 3.

### 3.4 Cross-series intron recurrence

An exact donor–acceptor pair observed in two or more independent GEO series provides a check of cross-dataset reproducibility, but recurrence alone does not establish biological validity or exclude systematic mapping or reference-index artifacts. Of the GSE221720 loci, 209/464 (45%) share an intron with at least one other series. Restricting the comparison to the other two polyA libraries yields 179/464 (39%), and overlap with the Ribo-Zero fracture series alone is 119/464 (26%). GSE193335 is the low outlier at 98/594 (17%), consistent with its thinner residual support (mean treatment junction sum 13.1 versus 32.6, 23.5, and 22.3) and lower unique depth (8.3 million versus 17.3, 24.1, and 19.4 million).

The recurrence checks that follow switch units from loci (209/464; Section 3.3) to primary-chromosome introns (595 for GSE221720, which exceeds 464 because a single locus can contribute more than one intron); the two counts are not interchangeable. A placement null (200 draws) yielded a mean of 0 and a maximum of 0 overlapping GSE221720 primary introns, compared with 249 observed. A more stringent null employs the 93,457 unique leak coordinates that are not among the 626 residual intron coordinates: 69 appear in the residual table of another series (0.074%), compared with 249/595 (42%) for residual-to-residual recurrence. Because those excluded junctions are enriched for non-canonical and short introns, this contrast should be interpreted as an upper bound (**Figure 4**).

**Figure 4.**
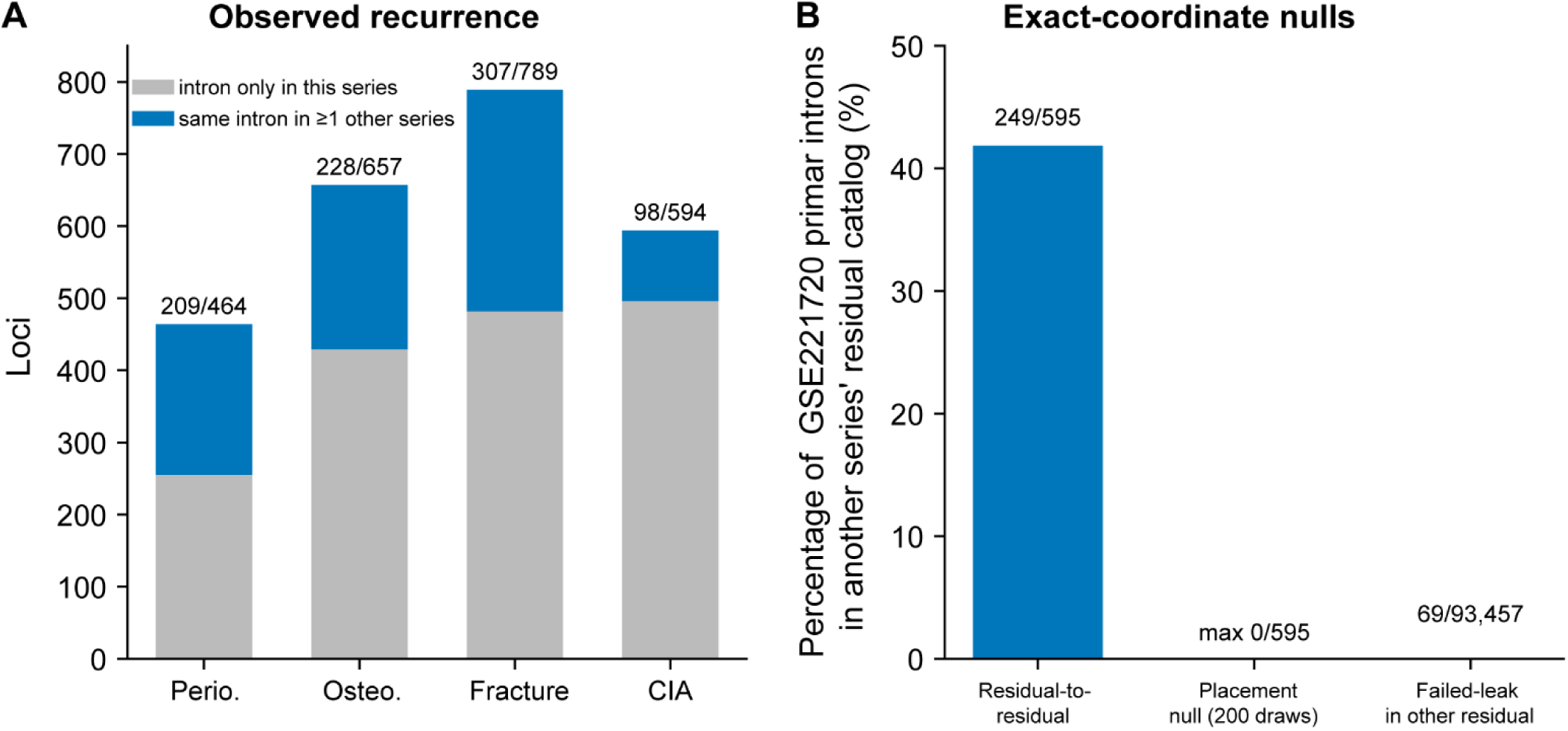
Cross-series intron recurrence. (**A**) Residual loci with an intron unique to that series versus present in at least one other series. (**B**) GSE221720 primary-chromosome introns (595) recurring in another series’ residual catalog: observed rate, the placement null (200 draws, max 0), and the failed-leak null (junctions that never entered the residual table).

Thirty-eight intron intervals are present in all four residual catalogs. Eighteen lie on unplaced contigs (GL456233.2, JH584295.1, and MU069434.1) and serve as a mapping-class positive control: the same STAR index can reproduce them from any BAM, and they are not reported as biological hits. The remaining twenty primary-chromosome intervals span seventeen residual loci (RSDL.23 and RSDL.30 each contribute more than one four-way intron) and are listed in Table 4. The three with a RepeatMasker fraction of 0 are RSDL.57, RSDL.137, and RSDL.168. The column nearest_distance_bp denotes the same-strand distance of the residual locus—the interval evaluated by the 1 kb structure gate. RSDL.137 lies 0.6 kb from Gm75657 and RSDL.168 abuts Efcab5 (0 bp); consequently, both fail that gate and remain four-way harvest hits rather than structure-pass intergenic models. RSDL.57, at 2.3 kb from Gm74130, is the only repeat-poor four-way primary interval that also passes the 1 kb gate.

**Table 4.** Primary-chromosome introns present in all four residual catalogs. . Donor and acceptor sites are GT-AG on the transcript strand. PAS denotes AATAAA or ATTAAA within the terminal 60 nt of the model. The column nearest_distance_bp is measured on the clipped locus—the interval used by the 1 kb structure gate—whereas the 200 nt harvest cutoff is evaluated on the intron span; a locus can therefore abut its nearest gene after terminal extension (0 bp) yet remain in the harvest catalog when its intron lies farther away. Three rows show 0 bp: RSDL.29 (Gm56686, intron span 1.4 kb) and RSDL.157 (Gm8424, intron span 1.4 kb) are repeat-rich mapping-class hits, whereas RSDL.168 (Efcab5, intron span 0.3 kb) is not. All three, together with RSDL.137 at 0.6 kb from Gm75657, fail the 1 kb structure gate on clipped-locus distance despite passing the 200 nt harvest cutoff on intron span.

| Locus | Shared intron | Nearest (kb) | n_j | nt | rmsk | PAS |
| --- | --- | --- | --- | --- | --- | --- |
| RSDL.29 | chrX:162757498–162757985 (+) | Gm56686 (0.0) | 3 | 2885 | 79% | no |
| RSDL.157 | chrX:149705136–149706200 (–) | Gm8424 (0.0) | 1 | 3044 | 96% | no |
| RSDL.168 | chr11:77080083–77089799 (–) | Efcab5 (0.0) | 1 | 373 | 0% | no |
| RSDL.137 | chr2:29748739–29757618 (+) | Gm75657 (0.6) | 1 | 649 | 0% | no |
| RSDL.156 | chr16:19057196–19058205 (+) | Gm56600 (1.6) | 1 | 1635 | 100% | no |
| RSDL.23 | chr11:31657463–31657885 (–) +1 | Gm53678 (1.8) | 4 | 6844 | 48% | no |
| RSDL.57 | chr11:83631684–83634777 (+) | Gm74130 (2.3) | 2 | 262 | 0% | no |
| RSDL.59 | chr1:173652401–173653590 (–) | Gm7897 (2.9) | 2 | 373 | 12% | no |
| RSDL.139 | chr9:110541422–110545336 (–) | Pth1r (3.4) | 1 | 2747 | 28% | no |
| RSDL.155 | chr12:103414265–103417077 (+) | Ifi27 (4.5) | 1 | 2080 | 78% | no |
| RSDL.33 | chrX:134881589–134883128 (+) | Gm15016 (5.0) | 3 | 1649 | 63% | no |
| RSDL.147 | chr2:84322323–84324006 (–) | Gm13710 (5.1) | 1 | 3761 | 44% | no |
| RSDL.85 | chr3:30895897–30901016 (–) | Gm42199 (6.1) | 2 | 2778 | 89% | no |
| RSDL.37 | chr3:6305699–6308345 (+) | Gm46796 (7.9) | 3 | 2163 | 96% | no |
| RSDL.30 | chr4:11178675–11179877 (–) +2 | Ints8 (12.4) | 3 | 4398 | 98% | no |
| RSDL.68 | chr11:19976263–19977633 (–) | Actr2 (33.5) | 2 | 1074 | 51% | no |
| RSDL.2 | chr6:22523135–22523678 (+) | Gm67965 (69.9) | 13 | 2802 | 27% | no |

Fourteen of the seventeen loci in **Table 4** have a RepeatMasker fraction ≥ 0.1. The most recurrent primary intervals are therefore repeat-rich, which is the expected outcome if four-way recurrence is interpreted as a shared-index certificate rather than as a gene call. Across the harvest set, loci at ≥ 0.1 recur at 173/377 (46%) and loci below 0.1 at 36/87 (41%); thus, recurrence is, if anything, slightly more common among repeat-rich intervals. The three four-way primary intervals below 0.1 are RSDL.57, RSDL.137, and RSDL.168. Taken together, the four-way overlap set—eighteen unplaced-contig introns plus fourteen of the seventeen repeat-rich primary loci—is better interpreted as evidence for a shared mapping and index artifact class than for shared transcription. RSDL.57 is the only repeat-poor four-way primary interval that also passes the structure gates; RSDL.137 and RSDL.168 fail the 1 kb gate at 0.6 kb from Gm75657 and 0 kb from Efcab5, respectively. BAM coverage for RSDL.57 and RSDL.137, and for the three contrast-screen rows, is shown in Figure 5.

**Figure 5.**
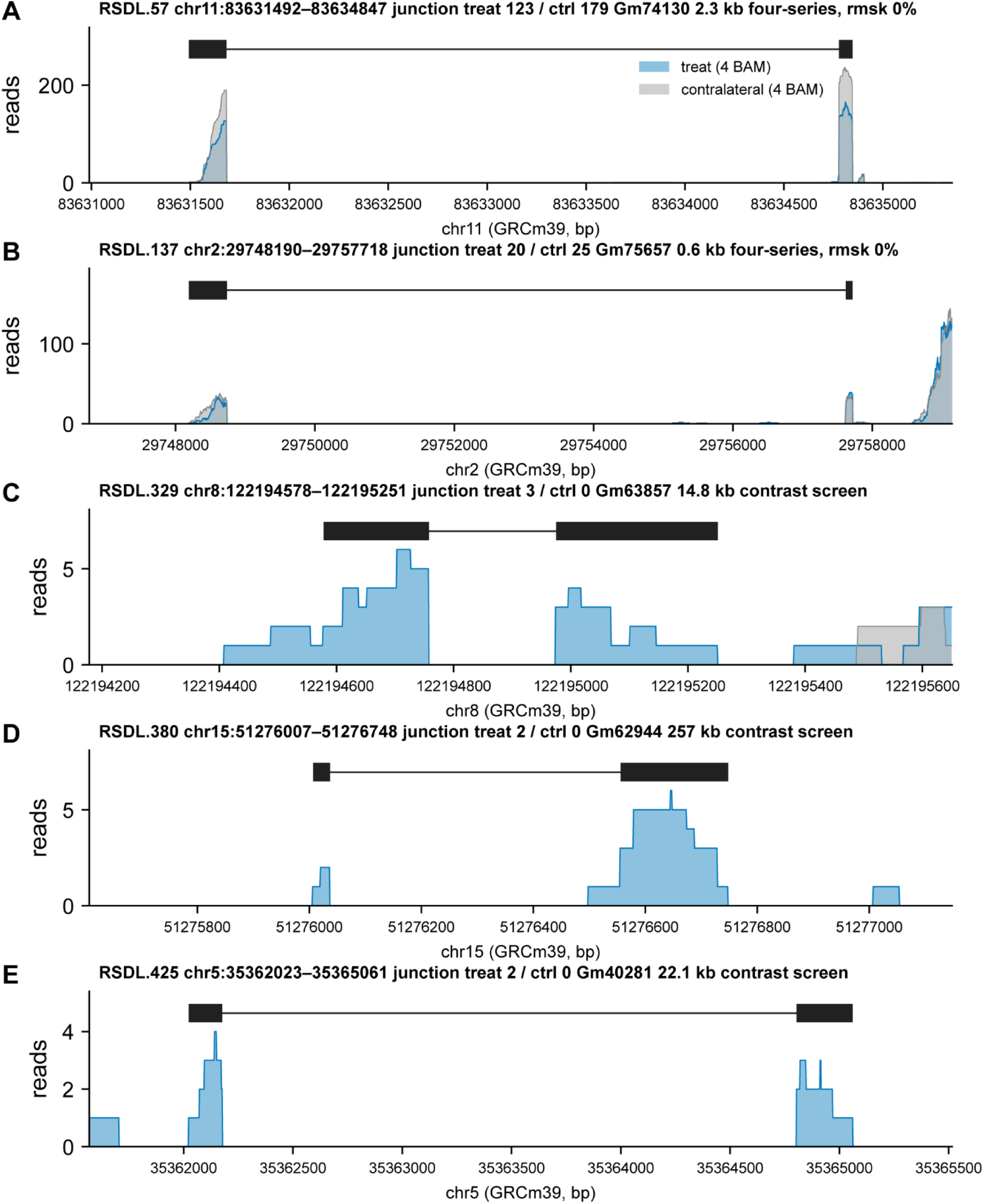
BAM coverage and junction-spanning reads on GSE221720. Tracks represent library sums rather than per-sample profiles. (**A, B**) Four-series repeat-poor loci RSDL.57 and RSDL.137, shown to illustrate boundary behavior at the 1 kb gate rather than as screen candidates: RSDL.57 passes the structure gates, whereas RSDL.137 fails on distance (0.6 kb) and is a harvest-only hit; junctions are present in treatment and contralateral libraries for both. (**C–E**) Contrast-screen rows, with two or three treatment fragments in total and no contralateral junction. Blue, sum of four treatment BAM files; gray, four contralateral BAMs. Black bars denote residual exons.

PAS hexamers (AATAAA or ATTAAA) occur within the terminal 60 nt of 53/464 GSE221720 residual models (11.4%; **Figure 6a**). By two-sided Fisher exact test, this rate does not differ significantly from that of 400 random genomic 60-mers (32/400, 8.0%; P = 0.11), yet it remains far below the rates at GENCODE protein-coding 3′ ends (253/400, 63%; P = 9 × 10⁻⁶⁰) and lncRNA 3′ ends (232/400, 58%; P = 1 × 10⁻⁴⁹). The more stringent recurrence null, comparing residual-to-residual recurrence with the failed-leak set, is shown in **Figure 4B**. This pattern is not an empty-walk artifact: only 1/464 3′ exons are 30 nt stubs, and 317/464 3′ exons exceed 200 nt. In this comparison, residual terminals therefore do not show a significant enrichment over random sequence and differ strongly from annotated 3′ ends. The coverage walk does not search for a PAS, and a long 3′ exon is not evidence of a mature polyadenylated gene. A score > 0 on the hexamer log-likelihood ratio is not a coding call (specificity 47.6% on GENCODE lncRNA).

**Figure 6.**
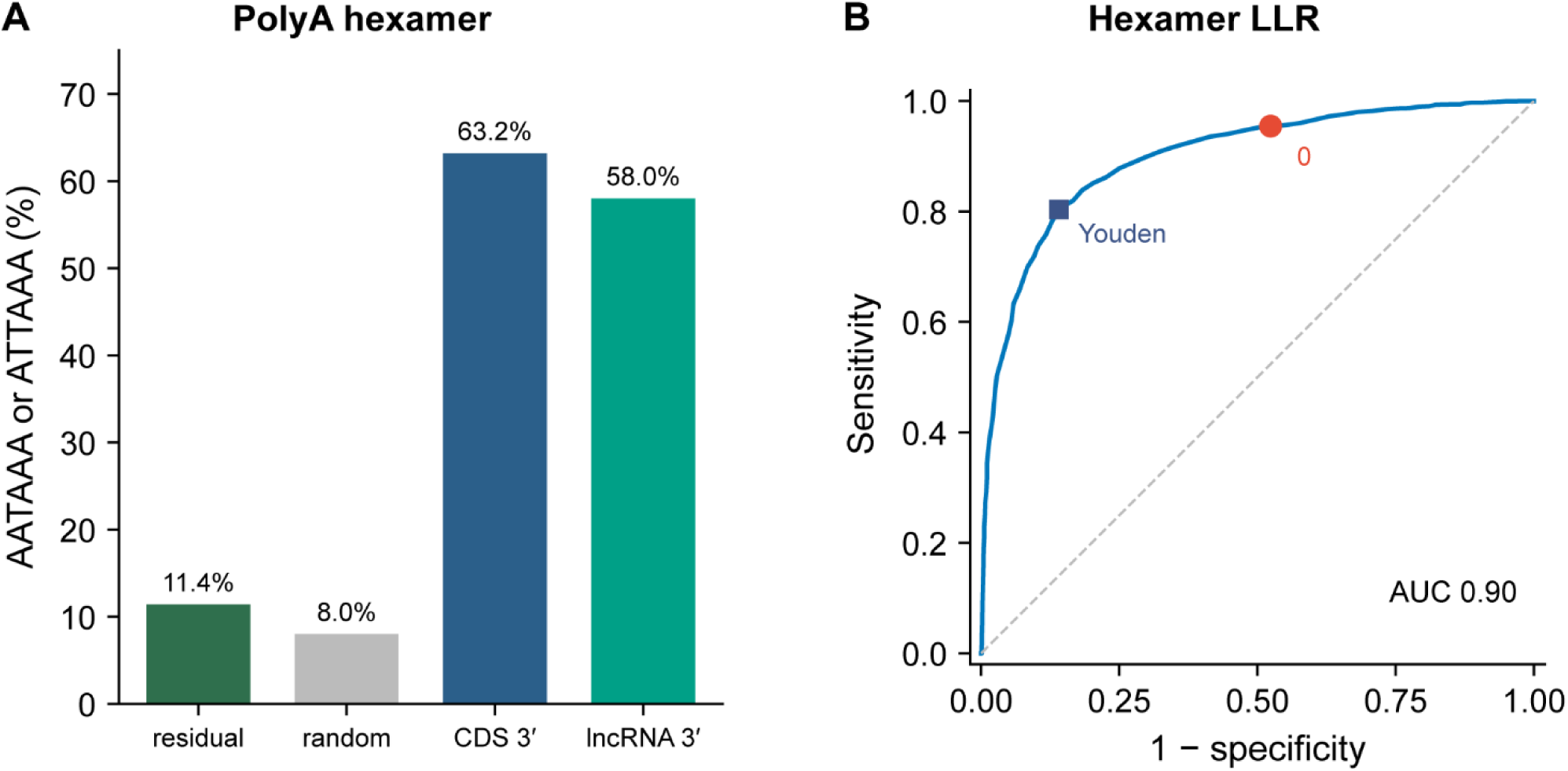
Sequence-based orthogonal checks. (**A**) PAS hexamer rate within the terminal 60 nt of residual models, in random genomic sequence, and at GENCODE 3′ ends. (**B**) ROC of the packaged hexamer LLR on GENCODE M39 protein-coding versus lncRNA transcripts; the circle marks the product threshold of 0, and the square marks Youden’s J.

**Figure 7.**
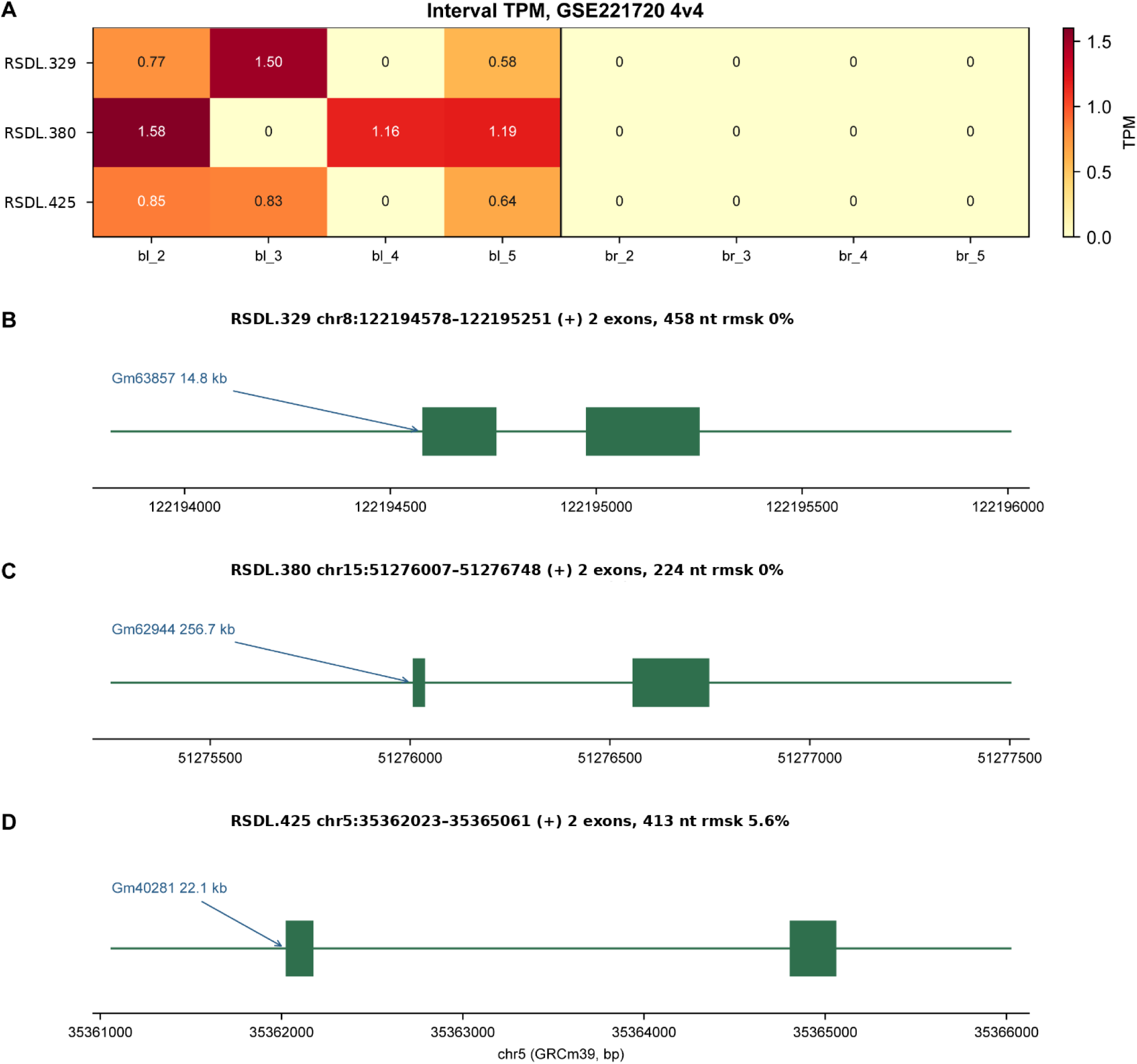
Contrast screen on GSE221720. (a) Interval TPM. Samples bl_2–bl_5 represent ligatured bone; br_2–br_5 represent unligated contralateral bone. (b–d) Exon maps.

Scored on the same GENCODE mRNA and lncRNA transcripts, the hexamer log-likelihood ratio of Section 2.7 achieves an AUC of 0.90 (Figure 6b). At the product threshold of 0, 95% of mRNA and 52% of lncRNA transcripts score positive. Residual-model scores overlap the lncRNA distribution: 97 of 227 scored ORFs exceed 0, with a median of −0.041, compared with 0.40 for mRNA and 0.013 for lncRNA.

### 3.5 Workflow demonstration: optional contrast filter

This section demonstrates the optional contrast filter and does not constitute a positive discovery. The four unligated contralateral BAM files were included as controls. Structure-pass residual loci were separated by interval TPM (Section 2.5): control maximum TPM < 0.5, detection in at least three treatment samples, treatment median TPM ≥ 0.5, and a RepeatMasker fraction < 0.1. Loci with shared harvest junctions were removed from the final table.

Under those gates, three rows remain (Table 5), all two-exon models of 224–458 nt. Of the C(8,4) = 70 ways to label four of the eight libraries as treatment, one is the observed assignment, twenty (lp01–lp20) were used to set the gates and are not reused, and the remaining 49 were never examined during gate setting. Across those 49 draws the mean is n = 0.24 (41/49 empty; maximum 2), and none reached n ≥ 3, the observed count, which bounds the permutation P at (0 + 1)/(49 + 1) = 1/50 = 0.02. The empirical size false discovery rate (FDR) on the same hold-out is approximately 0.08. These 49 draws constitute the complete relabeling of a single set of eight libraries rather than independent samples; consequently, the individual counts are correlated by construction. The tail probability remains the standard exact-permutation reference statistic; however, because the gate thresholds were fixed using labels drawn from that same eight-library pool, both quantities should be regarded as optimistic rather than conservative. Harvest was not re-run, so this is a gate-size FDR rather than a discovery FDR. The three rows are not Wald-significant, they share no intron with the other catalogs, and Figure 5 shows that their junction support amounts to two or three treatment fragments in total.

**Table 5.** Contrast screen on GSE221720. . Treatment samples are ligatured bone (bl_2–bl_5); contralateral samples are unligated bone (br_2–br_5). Distance is to the nearest same-strand gene in GENCODE M39.

| Locus | Interval | nt | Nearest (kb) | rmsk | bl_2 | bl_3 | bl_4 | bl_5 | br | DE |
| --- | --- | --- | --- | --- | --- | --- | --- | --- | --- | --- |
| RSDL.329 | chr8:122194578–122195251 (+) | 458 | Gm63857 (14.8) | 0% | 0.77 | 1.50 | 0 | 0.58 | 0 | low_count |
| RSDL.380 | chr15:51276007–51276748 (+) | 224 | Gm62944 (256.7) | 0% | 1.58 | 0 | 1.16 | 1.19 | 0 | low_count |
| RSDL.425 | chr5:35362023–35365061 (+) | 413 | Gm40281 (22.1) | 5.6% | 0.85 | 0.83 | 0 | 0.64 | 0 | low_count |

## 4. Discussion

The results support three main conclusions. First, existing bulk RNA-seq alignments contain recurrent spliced junctions outside GENCODE gene bodies, and many are reproducible across independent series (Section 3.4); this reproducibility does not by itself distinguish biological transcription from systematic mapping or index effects. Second, those junctions can be indexed without transcriptome assembly and counted in the same matrix as annotated genes, yielding hundreds of loci per series. A reference-guided StringTie merge reports comparable numbers of class-u loci (454, 189, and 1,102; Section 3.2, Table 2), but these are products of transcriptome reconstruction rather than a targeted intergenic splice census: after the same structure and detection gates, 7, 4, and 9 remain, and none reaches a contrast table. This illustrates the different operating point of TxNova relative to a standard StringTie workflow. Third, the 3′-end and coding-score features of residual models are more similar to the lncRNA comparison set than to mature protein-coding mRNA (Section 3.4). These conclusions should be interpreted within three important limitations.

Public bulk RNA-seq alignments can retain spliced reads outside current GENCODE annotations ^2^. When analyses are restricted to annotated gene-level count matrices, those signals are not represented as separate rows. TxNova converts recurrent intergenic junction evidence into residual intervals that share a count matrix with the annotation—an approach closer to a junction census ^17,18,20^ than to transcriptome assembly ^5,19^.

Reference-guided StringTie 3.0.3 does emit novel models from these BAM files; however, after the same structure and detection gates only 7 MSTRG loci remain on GSE221720, and none reaches the final contrast table. Residual harvest instead begins from the unassembled junction remainder (Table 2).

The most closely related tools operate at the junction level yet pursue different goals. LeafCutter clusters intron excisions and tests for differential splicing; its input mixes annotated and unannotated junctions, and its output is a cluster-level P value rather than a countable interval model ^17^. Portcullis filters spurious alignment-derived junctions and returns junctions rather than loci ^18^. ASJA extracts linear, back-splice, and fusion junctions ^20^. JunctionSeq tests novel junction usage at annotated genes ^22^. intropolis inventories unannotated splices across public RNA-seq ^21^. PsiCLASS and StringTie assemble a complete transcriptome ^5,19^. TxNova occupies the space between these approaches: it retains only junctions that are absent from the annotation and recurrent across samples, clusters them into loci, and places those loci in the same count matrix as annotated genes. The cost is narrow coverage—intronic and antisense channels are excluded, and neither long-read assembly nor differential splicing of annotated junctions is addressed.

Three limitations bound what these catalogs can support. First, residual models are fragments whose terminals are coverage walks rather than RACE ends. On GSE221720, the 3′ exon already exceeds 200 nt at 317/464 loci, yet PAS content does not differ significantly from random sequence and hexamer scores overlap the lncRNA range. Figure 5 shows that the two repeat-poor four-series loci RSDL.57 and RSDL.137 carry recurrently aligned junctions in both treated and contralateral libraries; consequently, they are shared residual signals rather than evidence for treatment-induced genes. No RT-PCR or long-read confirmation is reported here. Long-read RNA-seq (PacBio Iso-Seq or ONT direct RNA/cDNA) could in principle resolve full-length residual transcript structure and observed 3′ ends in a single run, in place of the coverage-walk terminal employed here; testing that combination remains future work.

Second, catalog precision is not estimated. Masked-gene recall measures sensitivity only. The placement and failed-leak nulls show that residual-to-residual recurrence is substantially above the coordinate-based backgrounds tested here, but they do not establish biological identity. The decoy control (0/5,000) shows that unsupported intergenic GT-AG pairs of annotated intron length essentially never coincide with a treatment-recurrent unassembled junction under this placement test; however, because decoy coordinates carry no simulated read support to begin with, that null has limited power and constitutes a placement control rather than a read-level precision estimate. None of these results assigns a false- positive rate to the remaining rows of Table 3. A spike-in simulation—transcripts relocated to intergenic sequence, resequenced at empirical depth, and realigned—is the experiment that would place a number on precision, and it is not attempted here. Unplaced and repeat-rich four-way hits are mapping-class positives; RSDL.57 is the only repeat-poor four-way primary interval that is also structure-pass, whereas RSDL.137 and RSDL.168 fail the 1 kb gate.

Third, no wet-lab identity test is included, and the software does not call antisense or intronic residual channels—a deliberate scope decision that keeps this first release a clean intergenic census. In GSE221720, 93,451 of 94,348 leak junction records overlap a gene body, while 897 do not. Shared harvest labels predominate in every catalog, suggesting that these data support the presence of recurrent unannotated junctions in the libraries rather than injury-specific novel genes. Human data are supported by the command-line interface but are not demonstrated here; mouse results should not be assumed to transfer without independent benchmarking. The hexamer AUC of 0.90 describes mRNA versus lncRNA discrimination on GENCODE rather than residual classification, and a positive score does not constitute a coding call. A comprehensive GTF is required because a cellranger-thin or basic annotation causes real genes to appear intergenic in a manner that preflight cannot detect.

The intended use is to take existing BAM files, index the intergenic residual intervals that an annotation-restricted gene-level analysis does not represent as separate features, and then—when control BAM files are present on the sheet—report treatment-detected, control-silent rows as a short screen. Like early lincRNA cataloging efforts ^27,39^, these outputs are intended as ranked resources for follow-up rather than definitive gene lists.

## Software and data availability

Analyses in this paper use TxNova 0.1.8 (https://doi.org/10.5281/zenodo.21970482).

Source: https://github.com/leelieber2025/TxNova

Bioconda(https://anaconda.org/bioconda/txnova)

PyPI (https://pypi.org/project/txnova/)

Documentation: https://txnova.readthedocs.io/

License: Apache-2.0.

Public FASTQ data are available under accessions GSE221720, GSE166522, GSE157460, and GSE193335 (Section 3.1).

## Ethics

This study reanalyzes publicly deposited mouse RNA-seq data. No new animal procedures were performed.

## Author contributions

Z.L. conceived the method, implemented the software, designed and performed the analyses, and wrote the manuscript. A.W.J. edited the manuscript. SX.L. implemented the software.

## Acknowledgements

We thank the investigators who deposited GSE221720, GSE166522, GSE157460, and GSE193335.

